# Bidirectional sex-dependent regulation of α6 and β3 nicotinic acetylcholine receptors by protein kinase Cε

**DOI:** 10.1101/2020.02.27.966978

**Authors:** Janna K. Moen, Margot C. DeBaker, Julia E. Myjak, Kevin Wickman, Anna M. Lee

## Abstract

Nicotine and alcohol are the most commonly abused substances worldwide, and comorbid nicotine and alcohol addiction is highly prevalent. Nicotinic acetylcholine receptors (nAChRs) containing the α6 and β3 subunits are expressed in neural reward circuits and are critical for both nicotine and alcohol reward. nAChRs are dynamically regulated by signaling molecules such as protein kinase C epsilon (PKCε), which impact transcription of α6 and β3 subunit mRNA (*Chrna6* and *Chrnb3*, respectively). Previous work found decreased expression of *Chrna6* and *Chrnb3* transcripts in the ventral midbrain of male PKCε^−/−^ mice, who also consume less nicotine and alcohol compared to wild-type (WT) littermates. Here, we show that female PKCε^−/−^ mice have enhanced expression of *Chrna6* and *Chrnb3* transcripts in the ventral midbrain, which functionally impacts nAChR-dependent behavior, as female but not male PKCε^−/−^ mice exhibit locomotor hypersensitivity to nicotine. Female PKCε^−/−^ mice show no differences in alcohol-induced sedation compared to WT littermates, while male PKCε^−/−^ have enhanced sedation compared to WT mice, a phenotype that has previously been reported in α6^−/−^ mice. Female PKCε^−/−^ mice also show reduced depression-like behavior in response to systemic injections of varenicline compared to WT littermates, and this effect was absent in male mice. Additionally, we found that female PKCε^−/−^ mice show altered alcohol and nicotine consumption patterns in chronic voluntary two bottle choice assays. Our data reveal a bidirectional effect of sex in the transcriptional regulation of nicotinic receptors by PKCε, and highlight the importance of studying both sexes in preclinical animal models.

## Introduction

Nicotine and alcohol are the two most commonly abused drugs, and co-abuse of both substances is highly prevalent.^1,2^ Despite the high costs associated with alcohol and nicotine dependence, current pharmacological approaches to assist cessation have limited success rates in achieving and maintaining abstinence.^3^ Additionally, sex influences clinical outcomes related to nicotine and alcohol abuse: women show a faster progression to the onset of alcohol use disorders and are less likely to seek treatment than men,^4^ and experience more difficulty when attempting to quit smoking.^5^ However, the majority of preclinical studies utilize only male animals, hindering our understanding of the neurobiological mechanisms underlying sex differences in alcohol and nicotine addiction.

Nicotinic acetylcholine receptors (nAChRs) are widely expressed pentameric ion channels that are implicated in both alcohol and nicotine reward mechanisms. Nicotine acts as a potent and selective agonist at neuronal nAChRs, while alcohol potentiates nAChR signaling through allosteric modulation.^6^ nAChRs are found presynaptically and on neuronal cell bodies, and as such are poised to directly influence cell excitability and neurotransmitter release. nAChRs are also incredibly diverse, with 11 different subunits (α2-7, α9-10, β2-4) expressed in the mammalian brain that can form a variety of channels with distinct pharmacological properties.^7^ The nAChR system is involved in a variety of affective behaviors including reward, anxiety, and depressive-like phenotypes.

The high affinity α6 subunit is frequently co-expressed with the ancillary β3 subunit in a nAChR subtype denoted α6β3* (* indicating presence of additional subunits in the heteropentamer). These subunits are preferentially expressed in dopamine neurons in the ventral midbrain and are critically involved in nicotine and alcohol reward, as mice lacking the α6 subunit (α6^−/−^ mice) fail to self-administer nicotine,^8^ and drugs targeting the α6 subunit can reduce alcohol and nicotine consumption in rodents.^9–11^ The genes encoding the α6 and β3 nAChR subunits (*Chrna6* and *Chrnb3*, respectively) are located back-to-back on chromosomes 8 in both humans and mice in what is termed the CHRNB3-CHRNA6 gene cluster. Small nucleotide polymorphisms in this region are associated with both alcohol and nicotine dependence in humans, further implicating α6β3* nAChRs in alcohol and nicotine reward mechanisms.^12,13^ Due to their close chromosomal proximity and colocalization *in vivo*, these genes are likely coregulated, but the transcriptional mechanisms underlying regulation of the CHRNB3-CHRNA6 cluster remain largely unknown.

One potential regulator of the CHRNB3-CHRNA6 cluster is the signaling molecule protein kinase C epsilon (PKCε). PKCε, a member of the serine/threonine kinase family, interacts with α4β2-containing nAChRs via phosphorylation to assist recovery from desensitization.^14^ Male mice lacking PKCε (PKCε^−/−^) also show reduced expression of *Chrna6* and *Chrnb3* in the ventral midbrain (VMB) and striatum, suggesting the involvement of PKCε in transcriptional regulation of this gene cluster.^15^ Further, male PKCε^−/−^ mice consume less alcohol and nicotine compared to their wild-type (WT) littermates, and show enhanced sensitivity to the sedative properties of alcohol.^15,16^ Recent studies have shown that targeting PKCε with small molecule inhibitors or conditional genetic inhibition can reduce alcohol consumption in mice, indicating that PKCε may be a promising target for pharmacotherapies to reduce alcohol and/or nicotine consumption.^17,18^ However, the majority of studies on the relationship between PKCε, nAChRs, and addictive behaviors have used only male mice, hindering the ability to translate preclinical research into a clinical setting.

The goal of the current study was to determine whether PKCε similarly influences nAChR subunit expression and associated behaviors in female mice. In contrast to prior work showing a decrease in *Chrna6* and *Chrnb3* transcript in the VMB of male PKCε^−/−^ mice compared to WT littermates,^15^ we found that female PKCε^−/−^ mice show higher expression of *Chrna6* and *Chrnb3* transcripts in the VMB compared to female WT littermates. This increase in transcript level is associated with enhanced sensitivity to the locomotor stimulatory effect of nicotine. While both male PKCε^−/−^ and α6^−/−^ mice show enhanced alcohol-induced sedation,^10,16^ we show that female PKCε^−/−^ mice have similar sensitivity to alcohol-induced sedation as their WT littermates. Additionally, female PKCε^−/−^ mice exhibited reduced depressive-like behavior in response to the nAChR partial agonist varenicline compared to WT littermates, an effect that was absent in male PKCε^−/−^ mice. Further, α6 and β3 nAChR upregulation in female PKCε^−/−^ mice resulted in enhanced nicotine consumption during the first week of a 4-week 2-bottle choice assay compared to WT littermates, while female PKCε^−/−^ mice exhibit decreased alcohol consumption. Interestingly, there was no significant difference in α6-sensitive somatodendritic ACh currents between female WT and PKCε^−/−^ mice in putative VTA dopamine neurons measured using the antagonist α-conotoxin PIA. Taken together, our data reveal a bidirectional effect of sex in the transcriptional regulation of α6 and β3 nAChR subunits by PKCε, highlighting the importance of studying both sexes in preclinical animal models.

## Materials and Methods

### Animal subjects

PKCε knockout mice were generated as described in Khaser et al 1999,^19^ and breeding pairs were generously provided by Dr. Robert Messing (University of Texas Austin). The PKCε null allele was maintained on an inbred 129S4/SvJae background. PKCε^+/−^ 129S4/SvJae mice were crossed with C57BL6/J mice to generate PKCε^+/−^ C57BL6/J x 129S4/SvJae F1 hybrid breeding pairs, which were used to generate hybrid PKCε^−/−^ and wild-type (PKCε^*+/+*^, WT) littermates for experiments. Male and female mice for slice electrophysiology were 45-55 days old on the day of experiment, all other mice were a minimum of 56 days old. Mice were housed in a 12 hr light/dark cycle room in groups of 5 or fewer per cage with food and water provided ad libitum. Animals were group housed for all experiments unless otherwise indicated. Female animals were freely cycling and not monitored for estrous cycle state. All procedures were conducted under guidelines established by the University of Minnesota Institutional Animal Care and Use Committee and conformed to NIH guidelines.

### Drugs

For consumption studies, alcohol (Decon Labs, King of Prussia, PA) or nicotine tartrate (Acros Organics, Thermo Fisher Scientific, Chicago, IL) were mixed with tap water to the concentrations reported for each experiment. Nicotine concentrations are reported as free base, and nicotine solutions for consumption studies were not filtered or pH adjusted. Nicotine solution was masked with saccharin (Sigma Aldrich, Saint Louis, MO). Varenicline tartrate and desipramine hydrochloride were commercially purchased (Tocris Biosiences, Bio-Techne, Minneapolis, MN) and dissolved in 0.9% saline and were not filtered or pH adjusted. Injectable nicotine tartrate was dissolved in 0.9% saline to a stock concentration of 1.0 mg/mL and adjusted to pH 7.4. Stock solution was diluted to 0.25 mg/mL in 0.9% saline for a standard injection volume of 10.0 mL/kg. Injectable alcohol was dissolved in 0.9% saline for a final concentration of 20% v/v and was not pH adjusted.

### Tissue qPCR

Drug-naïve mice were deeply anesthetized with isoflurane followed by decapitation. Brains were rapidly dissected over ice into components from the striatum, ventral midbrain, and amygdala. To extract RNA, samples were homogenized via mortar and pestle followed by needle aspiration, then prepared according to QIAGEN RNeasy Plus kit (QIAGEN, Germantown, MD). RNA quality and concentration were measured using a Nanodrop 2000c spectrophotometer (Thermo Fisher Scientific, Chicago, IL). 500-1000 ng RNA per sample was reverse-transcribed using Applied Biosystems high-capacity reverse transcription kit (Applied Biosystems, Foster City, CA). Quantitative RT-PCR was performed using Taqman probes commercially available from Applied Biosystems targeting GAPDH (Mm99999915_g1) and *Chrna4* (Mm00516561_m1), *Chrna6* (Mm00517529_m1), *Chrnb2* (Mm00515323_m1), *Chrnb3* (Mm00532602_m1), and *Prcke* (Mm00440894_m1). Expression levels relative to GAPDH were calculated using the double-deltaCT method, with WT deltaCT levels averaged for a relative expression value of 1.

### Nicotine-induced locomotor activity

The nicotine-induced locomotor activity assay was adapted from Drenan et al, 2008.^20^ Drug-naïve mice were handled and acclimated to saline injections (10.0 mL/kg *i.p.*) for 2 days. On day 1, mice were placed in an open locomotor activity chamber (MedAssociates, Fairfax, VT) for 30 minutes to assess spontaneous locomotion in a novel environment. On day 2, mice were placed into the open locomotor activity channel. After 8 minutes, mice were removed, injected with 0.25 mg/kg nicotine *i.p.* or equivalent volume of saline, and placed back in the chamber within 15 seconds. Mice remained in the chamber for an additional 32 minutes for a total session time of 40 minutes. Locomotor activity was measured by beam breaks (ambulatory counts). For statistical analyses, data were pooled into 5 minute bins, with bin 0=5 minutes post-injection.

### Loss of righting reflex (LORR)

LORR procedure was adapted from Crabbe et al, 2006.^21^ Drug-naïve mice were injected with 4.0 g/kg alcohol *i.p.* and placed in an individual cage, observed for signs of intoxication, and placed on their back. Righting reflex was considered lost when the animal was unable to right itself for at least 30 seconds within 3 minutes of injection. Animals that failed to display LORR within 3 minutes after injection were excluded from analysis due to high probability of an off-target injection, which has previously been correlated with low blood alcohol content.^21^ Righting reflex was considered regained when the animal could right itself 3× within 30 seconds.

### Tail suspension test

Drug-naïve mice were injected *i.p.* with varenicline (1.0 mg/kg), desipramine (20.0 mg/kg), or equivalent volume of saline (10.0 mL/kg) 30 minutes prior to testing. Tails were attached to PE20 tubing using standard laboratory tape and gently suspended for 6 minutes and recorded on video camera. Videos were scored by a blinded experimenter for total time spent immobile, with immobility defined as the lack of any movement except involuntary swinging after a bout of movement.

### Nicotine 2-bottle choice

Female drug-naïve mice were singly housed in double grommet cages. Procedure was carried out as described in Lee and Messing 2011.^15^ Briefly, mice were presented with a bottle of tap water with 2% w/v saccharin and a bottle of tap water containing 15 μg/mL nicotine and 2% w/v saccharin for 4 weeks. Bottles were weighed every 2-3 days, and the positions of the bottles were alternated to account for side preferences. Mice were weighed once a week throughout the study. Evaporation and spillage were accounted for using control bottles in an empty cage.

### Alcohol 2-bottle choice

The alcohol 2-bottle choice assay was carried out as previously described by our group.^22,23^ Briefly, drug-naïve mice were single housed and presented with a bottle of tap water and a bottle of tap water containing increasing concentrations of alcohol: 3%, 6%, 10%, 14%, and 20% v/v. Each concentration was presented for 4 days. Bottles were weighed every 2 days, and the positions of the bottles were alternated to account for side preferences. Mice were weighed once a week throughout the study. Evaporation and spillage were accounted for using control bottles in an empty cage.

### Preparation of brain slices and patch clamp electrophysiology

Horizontal slices (225 μm) containing the VTA were prepared in ice-cold sucrose ACSF as described previously,^24^ and allowed to recover at room temperature in ASCF containing (in mM): 125 NaCl, 2.5 KCl, 1.25 NaH_2_PO_4_, 25 NaHCO_3_, 11 Glucose, 1 MgCl_2_, and 2 CaCl_2_, pH 7.4. for at least 1 hr. Neurons found in the lateral aspect of the VTA, just medial to the medial terminal nucleus of the accessory optic tract (MT), were targeted for recording. Whole-cell data were acquired using a Multiclamp 700B amplifier and pCLAMPv.9.2 software (Molecular Devices, Sunnyvale, CA). Putative DA neurons were selected based on morphology, size (apparent capacitance >40 pF), I_h_ current (>80 pA), and spontaneous activity (<5 Hz) (**table 1**), as these properties correlate well with tyrosine hydroxylase expression.^24^ The current response to a 1s voltage ramp (−60 to −120 mV) was used to assess I_h_ amplitude, and spontaneous activity was measured in current-clamp mode (I=0) for 1 min. In addition, the D2 DA receptor (D2R) agonist quinpirole (20 μM, Sigma, St. Louis, MO) was applied at the end of each experiment as a pharmacological test of DA neuron ientity.^25^ Whole-cell somatodendritic nAChR-mediated currents were evoked in the presence of tetrodotoxin (TTX) (0.5 μM, Tocris Bioscience, Bristol, UK) by locally applying acetylcholine (ACh; 1 mM, Sigma) via pressure ejection using a Picospritzer III (General Valve, Fairfield, NJ). The superfusion medium contained atropine (1 μM, Sigma) at all times to block muscarinic activity. α-conotoxin PIA (75 nM, Alomone Labs, Jerusalem, Israel) was bath applied for 12 minutes before local ACh application to measure the α6-insensitive component. The α6-sensitive component was calculated by taking the difference between baseline ACh currents and ACh currents recorded in the presence of α-conotoxin PIA, then dividing that by the baseline ACh current and multiplying by 100.

**Table 1.** Average whole-cell parameters from putative VTA dopamine neurons targeted for slice electrophysiology. Mean ± SEM.

|  | MALE |  | FEMALE |  |
| --- | --- | --- | --- | --- |
| Genotype | WT | PKCε<sup>-/-</sup> | WT | PKCε<sup>-/-</sup> |
| Capacitance (pF) | 59 ± 5 | 66 ± 5 | 63 ± 3 | 63 ± 6 |
| I<sub>h</sub> (pA) | -200 ± 32 | -348 ± 67 | -242 ± 71 | -261 ± 96 |
| Frequency (Hz) | 1.4 ± 0.4 | 1.7 ± 0.5 | 1.3 ± 0.2 | 1.3 ± 0.2 |

### Statistical analyses

All analyses were performed using Prism 8 (GraphPad, La Jolla, CA). Data were tested for normality and variance, and outliers were detected using Grubb’s test. Welch’s corrections were used in cases of unequal variance. Correlation coefficients for nAChR transcript levels were calculated using Pearson’s correlation. Comparison of data across time or across treatment groups with two dependent variables used 2-way ANOVA followed by Sidak’s multiple comparisons tests. Comparisons between groups with a single dependent variable used unpaired two-tailed *t*-tests.

## Results

### Female PKCε^−/−^ mice show elevated α6 and β3 nAChR mRNA in the ventral midbrain and locomotor hypersensitivity to nicotine

We performed quantitative real-time PCR on tissue from female WT and PKCε^−/−^ mice to determine the expression levels of nAChR subunit transcript between genotypes. We found that female PKCε^−/−^ mice displayed significantly higher levels of *Chrna6* (Welch’s unpaired *t*-test, *t*=3.526, df=17.64, \*\**p*=0.003) and *Chrnb3* (Welch’s unpaired *t*-test, *t*=3.837, df=17.68, \*\**p*=0.001) mRNA expression in the ventral midbrain (VMB) compared to WT littermates (**fig. 1A**), with no changes in expression observed for *Chrna4* or *Chrnb2* mRNA. No changes in expression where observed between genotypes across nAChR subtypes in the amygdala (**fig. 1B**) or striatum (**fig. 1C**). We then tested whether transcript expression in the VMB varied between male and female WT animals and found no differences in expression levels of *Chrna4*, *Chrna6*, *Chrnb2*, or *Chrnb3* transcript (**fig. 1D**). We further measured the expression of PKCε mRNA (*Prcke*) in the VMB and observed no differences between sexes. Notably, *Chrna6* and *Chrnb3* transcript levels were highly correlated across individual animals in both female WT (Pearson’s correlation, *r*=0.94, r^2^=0.906, *p*<0.0001, **fig. 1E**) and KO (Pearson’s correlation, *r*=0.956, r^2^=0.913, *p*<0.0001, **fig. 1F**) animals.

**Figure 1.**
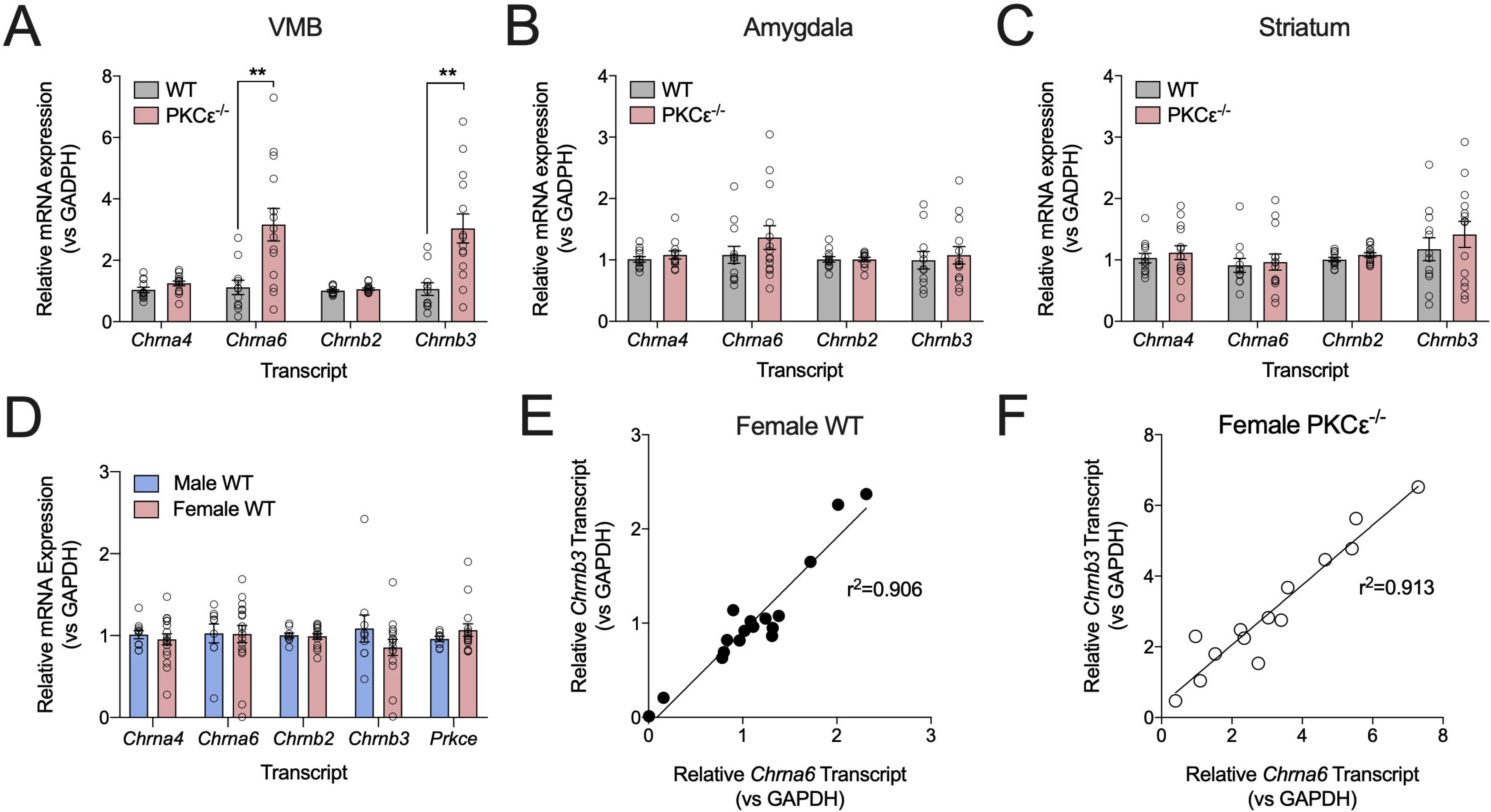
α6 and β3 nAChR mRNA is upregulated in the ventral midbrain of female PKCε^−/−^ mice. (**A**) A 3-fold increase of *Chrna6* and *Chrnb3* expression levels was observed in the ventral midbrain of female PKCε^−/−^ mice compared to WT littermates, with no differences in *Chrna4* or *Chrnb2* levels. No differences in transcript levels were observed between genotypes in the amygdala **(B)** or striatum **(C)**. Data shown as mean ± SEM, *n*=12-14 mice/genotype. \*\**p*<0.0025, unpaired t-test with Welch’s correction. **(D)** No differences were observed in endogenous expression of *Chrna4*, *Chrna6*, *Chrnb2*, *Chrnb3*, or *Prcke* transcript in the ventral midbrain of wild-type male and female mice. *n*=9-17/sex. **(E, F)** Transcript levels of *Chrna6* and *Chrnb3* are highly correlated across individual female WT **(E)** and PKCε^−/−^ **(F)** mice. Pearson’s r=0.9517 (WT, *p*<0.0001), =0.9555 (PKCε^−/−^, *p*<0.0001).

In order to determine whether elevated levels of α6 and β3 nAChR subunit mRNA impacted α6* nAChR-dependent behaviors, mice were injected with a low dose of nicotine (0.25 mg/kg *i.p.*) and monitored for locomotor activity (ambulatory counts). Female PKCε^−/−^ mice displayed a marked increase in ambulatory counts compared to female WT mice (**fig. 2A**). For statistical analysis, ambulatory counts were pooled into 5 minute bins, with bin 0=5 minutes post-injection (**fig. 2B**). In female mice, there was a significant genotype X time interaction as well as a significant main effect of both genotype and time (2-way RM ANOVA: F_genotypeXtime_(5,65)=4.592, *p*=0.001; F_genotype_(1,13)=5.100, *p*=0.04; F_time_(5,65)=18.2, *p*<0.001) with Sidak’s multiple comparisons revealing a significant genotype difference at bin 0 (**fig. 2B**). This was not due to an overall increase in locomotor activity, as there were no differences observed in ambulatory counts in a novel environment (**fig. 2C**) or in drug-naïve mice injected with an equivalent volume of saline (**fig. 2D**).

**Figure 2.**
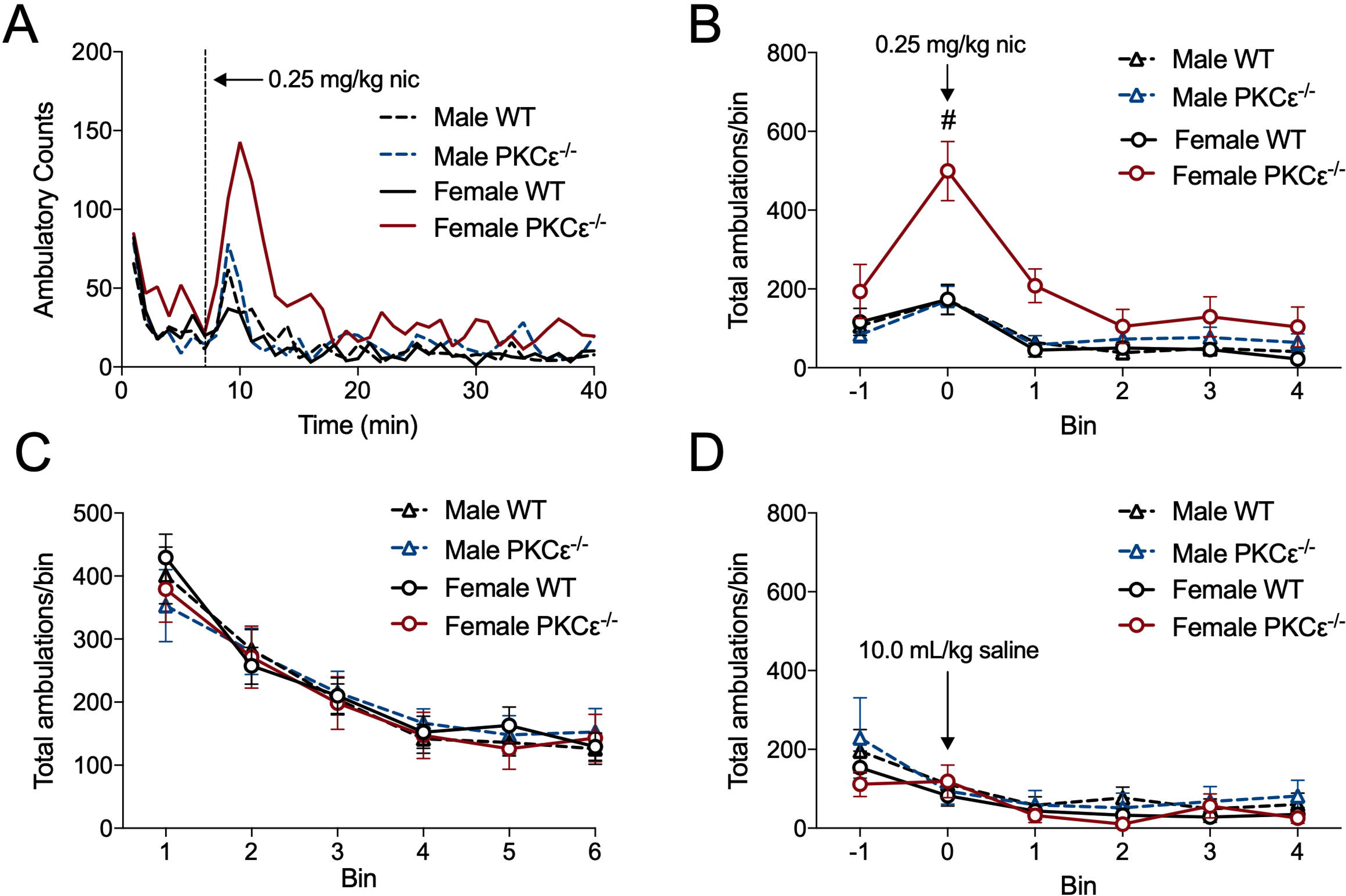
Female PKCε^−/−^ mice are hypersensitive to the locomotor stimulatory action of nicotine. (**A**) Average traces of ambulatory counts following 0.25 mg/kg nicotine injection. (**B**) Binned ambulatory counts (5 minute bin) show a significant effect of genotype in female mice. Sidak’s multiple comparisons reveal a significant difference (#) at bin 0. (**C**) no changes observed in baseline locomotor activity between sexes or genotypes. (**D**) Mice injected with 10.0 mL/kg saline show no changes in ambulatory activity. *n*=6-10 mice/group. Data shown as mean±SEM.

### Female PKCε^−/−^ mice show differences in nAChR-dependent acute behavior

We utilized the loss-of-righting reflex (LORR) assay to measure sensitivity to the sedative properties of alcohol across sexes and genotypes. We found that male PKCε^−/−^ mice showed increased alcohol-induced LORR duration compared to WT animals following an *i.p.* injection of 4.0 g/kg alcohol (unpaired 2-tailed *t*-test: t=5.261, df=12, \*\*\**p*=0.0002, **fig. 3A**), replicating prior findings in male PKCε^−/−^ mice.^16^ In contrast, we observed no difference in LORR duration between female WT and PKCε^−/−^ mice (unpaired 2-tailed *t*-test: t=1.052, df=19, *p*=0.31; **fig. 3B**).

**Figure 3.**
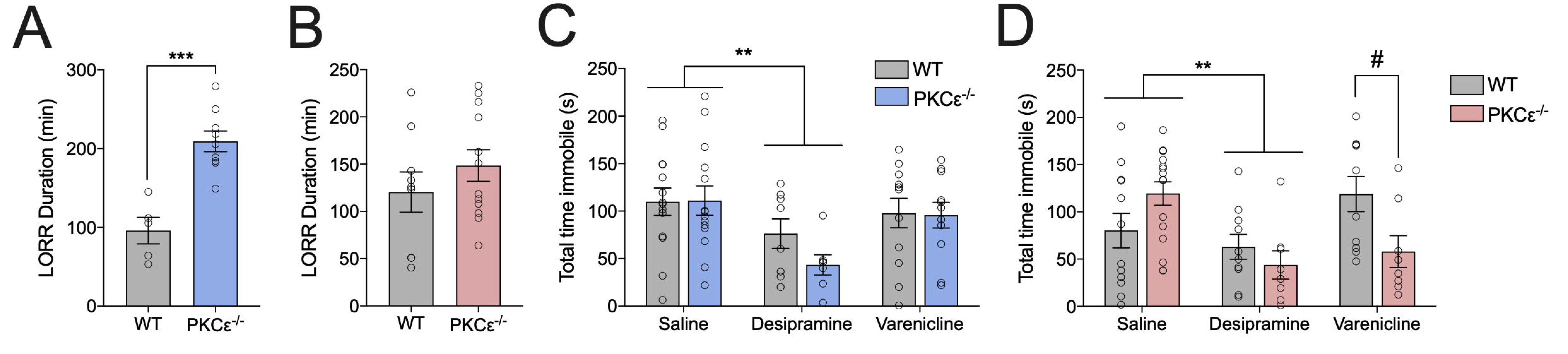
Differences in acute nAChR-dependent behavior between male and female PKCε^−/−^ mice. (**A**) Male PKCε^−/−^ mice show increased loss of righting reflex duration compared to WT animals following injection of 4.0 g/kg alcohol *i.p.* (unpaired two-tail *t*-test, \*\*\**p*=0.0002, *n*=5 WT, 9 KO). (**B**) Female mice show no differences in alcohol-induced sedation as measured by the loss of righting reflex. *n*=8 WT, 11 KO. (**C**) Male WT and PKCε^−/−^ mice show reduced immobility time in response to the tricyclic antidepressant desipramine in the tail suspension test (**Main effect of drug, *p*<0.005, *n*=7-14 mice/group). (**D**) Female WT and PKCε^−/−^ mice show reduced immobility time in response to desipramine (**Main effect of drug, *p*<0.005, 2-way ANOVA, *n*=8-16 mice/group). Female PKCε^−/−^ mice exhibit reduced immobility time in response to the nAChR partial agonist varenicline (#*p*<0.05 between genotypes by Sidak’s multiple comparisons test). Data shown as mean ± SEM.

In order to determine the impact of PKCε deletion on depression-like behavior, male and female WT and PKCε^−/−^ mice underwent the tail suspension test (TST) following injections of saline, the tricyclic antidepressant desipramine (20 mg/kg *i.p.*), and the nAChR partial agonist varenicline (1.0 mg/kg *i.p.*). As expected, desipramine reduced immobility time compared to saline in both males (2-way ANOVA: F_drug_(1,39)=9.92, \*\**p*=0.003; F_genotype_(1,39)=0.97, *p*=0.33; F_genotypeXdrug_(1,39)=1.12, *p*=0.30, **fig. 3C**) and females (2-way ANOVA: F_drug_(1,42)=9.921, \*\**p*=0.0031; F_genotype_(1,42)=0.42, *p*=0.52; F_genotypeXdrug_(1,42)=3.53, *p*=0.07, **fig. 3D**) regardless of genotype. While male mice showed no changes in immobility time in response to varenicline, there was a significant genotype × drug interaction in female mice (2-way ANOVA: F_genotypeXdrug_(1,42)=8.880, *p*=0.005; F_drug_(1,42)=0.4644, *p*=0.50; F_genotype_(1,42)=0.414, *p*=0.52, **fig. 3D**). Sidak’s multiple comparisons revealed that female PKCε^−/−^ mice showed a significant decrease in total time immobile following treatment with varenicline (**fig. 3D**).

### Chronic voluntary consumption of alcohol and nicotine in female WT and PKCε^−/−^ mice

Female WT and PKCε^−/−^ mice were presented with continuous access to a bottle of 15 μg/mL nicotine plus 2% w/v saccharin alongside a bottle of water and 2% w/v saccharin for 4 weeks. Repeated measures 2-way ANOVA indicated a genotype X time interaction for nicotine consumption (F_genotypeXtime_(3,189)=8.045; *p*<0.0001; F_time_(3,189)=0.273, *p*=0.72; F_genotype_(1,63)=0.15, *p*=0.70, **fig. 4A**), with Sidak’s multiple comparisons indicating that female PKCε^−/−^ mice had increased nicotine consumption during the first week. There were no significant changes in nicotine preference (2-way RM ANOVA: F_time_(3,189)=0.595, p=0.62; F_genotype_(1,63)=0.0002, p=0.97; F_timeXgenotype_(3,189)=1.376, p=0.25; **fig. 4B**).

**Figure 4.**
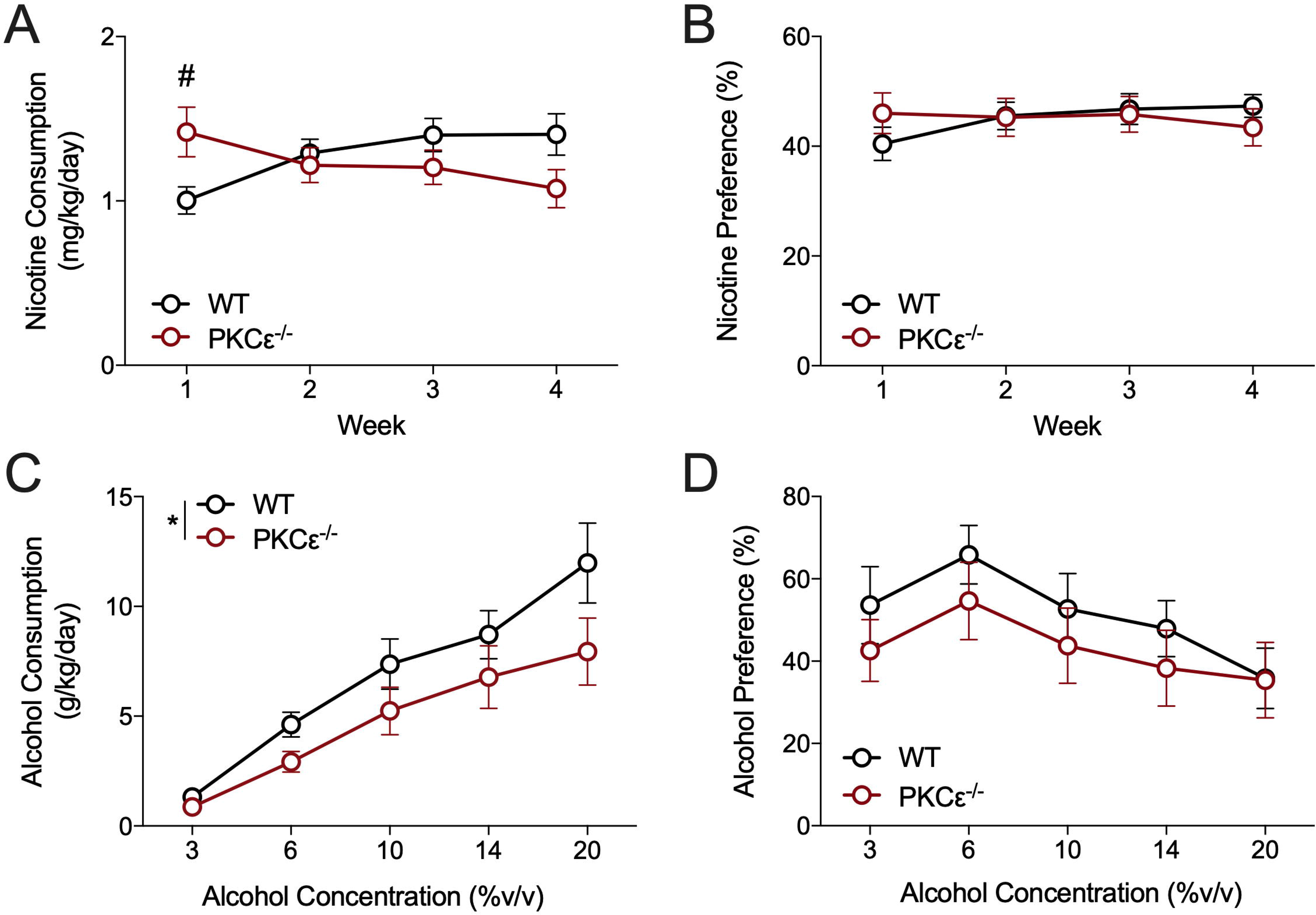
Female PKCε^−/−^ mice show distinct consumption patterns in chronic voluntary consumption assays. (**A**) Female PKCε^−/−^ mice consume more nicotine during the first week of a chronic voluntary nicotine two-bottle choice assay (#*p*<0.05 between genotypes by Sidak’s multiple comparisons test). (**B**) no genotype differences were observed in nicotine preference. *n*=30-35 mice/genotype. (**C**) Female PKCε^−/−^ mice show reduced consumption of alcohol compared to WT littermates (*Main effect of genotype, *p*<0.039 by RM 2-way ANOVA). (**D**) Female PKCε^−/−^ mice do not show any differences in alcohol preference compared to WT littermates. *n*=15/genotype. Data shown as mean±SEM.

We next performed a chronic, voluntary 2-bottle choice test in which female WT and PKCε^−/−^ mice were presented with 24 hour access to escalating concentrations of alcohol. Repeated measures 2-way ANOVA indicated a main effect of both concentration and genotype (2-way RM ANOVA: F_concentration_(4,112)=25.46, *p*<0.0001; F_genotype_(1,28)=4.682, \**p*=0.04; F_concentrationXgenotype_(4,112)=0.889, p=0.47; **fig. 4C**) on alcohol consumption. There was no main effect of genotype or any interaction between genotype and concentration on alcohol preference (2-way RM ANOVA: F_genotype_(1,28)=0.9507, *p*=0.34; F_concentration_(4,112)=3.668, *p*=0.008; F_concentrationXgenotype_(4,112)=0.227, p=0.92; **fig. 4D**).

### No sex or genotype differences observed in baseline or α6-sensitive ACh currents in VTA DA neurons

As α6 and β3-containing nAChRs are predominately expressed in DA neurons in the ventral midbrain,^26,27^ we used the α6 nAChR antagonist α-conotoxin PIA (α-CTX) to measure α6 nAChR-dependent somatodendritic ACh currents in putative VTA DA neurons using local ACh application (**fig. 5A**). No differences in baseline ACh currents were observed between sexes or genotypes (2-way ANOVA: F_sex_(1,57)=0.9357, p=0.33; F_genotype_(1,57)=3.067, p=0.09; F_genotypeXsex_(1,57)=0.267, p=0.60; **fig. 5B, C**). While we observe a trend towards a higher α6-sensitive component in female PKCε^−/−^ mice compared to female WT mice, this result was not statistically significant (unpaired 2-tailed *t*-test, *t*=1.104, df=15, p=0.29; **fig. 5E**).

**Figure 5.**
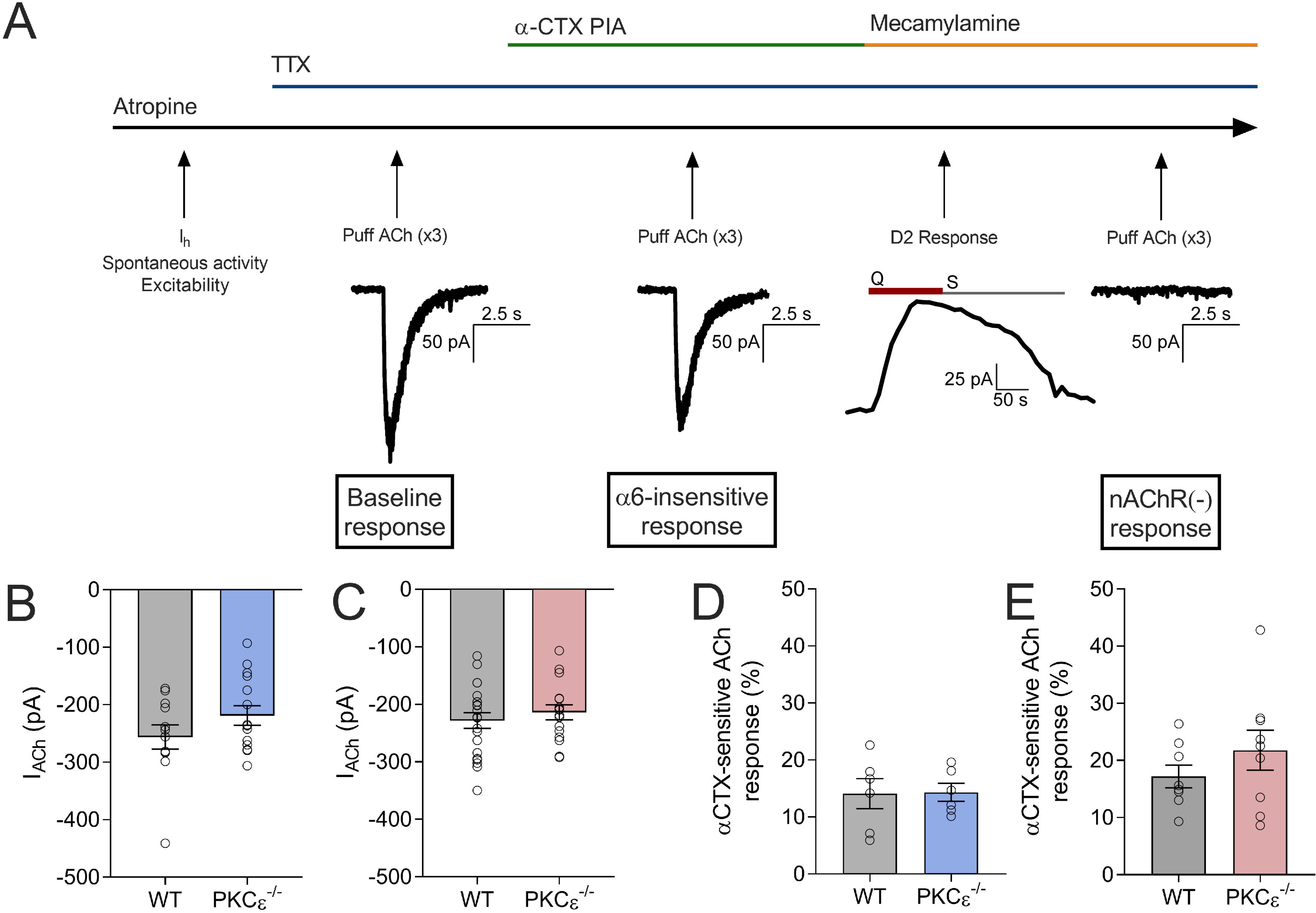
Measurement of α6-sensitive ACh currents in VTA dopamine neurons. (A) Experimental design for slice electrophysiology experiments. D2 response: Q=quinpirole, S=sulpiride. No differences were observed in baseline ACh currents in putative VTA DA neurons between genotypes in males (B) or females (C). α-CTX-sensitive ACh currents do not differ between genotypes in males (D) or females (E).

## Discussion

### Regulation of nicotinic acetylcholine receptors by PKCε

Previous work has shown that ablation of PKCε in male mice results in decreased expression of *Chrna6* and *Chrnb3* mRNA in the VMB.^15^ In contrast, we show elevated levels of *Chrna6* and *Chrnb3* mRNA in the VMB of female PKCε^−/−^ mice compared to WT littermates. We next asked whether baseline nAChR transcript levels differed between sexes and observed no changes in relative transcript expression between WT male and female mice. Notably, the expression of *Chrna6* and *Chrnb3* transcript was highly correlated across individual female WT and PKCε^−/−^ mice. Taken together with the close chromosomal proximity of these two genes in the CHRNB3-CHRNA6 gene cluster, our data suggest that they are coregulated in a process that involves PKCε signaling. As PKCε is a protein kinase that does not directly regulate gene expression, we speculate that PKCε is part of signaling cascade that results in changes to transcription factor activation or binding. However, to our knowledge, there have been no studies examining transcriptional regulatory elements or transcription factor binding sites surrounding the CHRNB3-CHRNA6 gene cluster, and the precise mechanism by which PKCε is able to impact transcript levels remains to be determined.

Despite our results showing changes in nAChR-associated behaviors, we did not observe a significant difference in α6-sensitive ACh currents in female PKCε^−/−^ mice. One major limitation of this experiment is that α-conotoxins, including α-CTX PIA, bind preferentially to the interface between α6 and β2 nAChR subunits.^28^ As such, α-CTX may not be an effective antagonist if α6 and β3 subunits are being incorporated into the same heteropentamer without an additional β2 subunit. While we speculate that α6 and β3 nAChR upregulation is occurring in dopamine neurons in the VTA due to their relatively selective expression profile,^26^ there are a variety of dopaminergic subpopulations that exhibit distinct inputs, projection targets, and cell markers.^29^ Additionally, α6* nAChRs have been observed at low levels in a subpopulation of VTA GABA neurons.^27^ The upregulation we observe using RT-qPCR may be limited to a neuronal subpopulation, a different nucleus within the VMB such as the substantia nigra pars compacta, or possibly result in preferential receptor expression in DA neuron terminals in the striatum. Additionally, receptor upregulation may impact nAChR function in a way that was not captured with our electrophysiological experiment, such as changes in receptor desensitization kinetics or agonist affinity.

### Influence of α6 nAChRs on acute behaviors

In order to determine whether the observed upregulation of nAChR transcript was correlated with behavioral consequences, we performed acute behavioral paradigms that have been shown to be influenced by α6* nAChRs. Mice engineered to express a hypersensitive α6 subunit exhibit enhanced locomotor activity in response to low doses of nicotine that are subthreshold in WT animals.^20^ Our data indicate that female PKCε^−/−^ mice are sensitive to the locomotor stimulatory action of low-dose nicotine, suggesting that increased α6 and β3 transcript expression resulted in relevant α6* nAChR-dependent behaviors. Importantly, the observed sensitivity to low-dose nicotine was not a feature of general hyperactivity, as locomotion upon being placed in a novel chamber environment was unaffected, and all mice showed gradual reduction of locomotor responses by the conclusion of the 30 minute test period.

Both α6^−/−^ and male PKCε^−/−^ mice exhibit enhanced alcohol-induced sedation as measured by the loss of righting reflex assay (LORR),^16,30^ indicating that the α6 nAChR subunit is an important contributor to the sedative properties of alcohol. We replicated the enhanced sensitivity to alcohol-induced sedation in male PKCε^−/−^ mice. In contrast, we found that female PKCε^−/−^ mice have similar LORR duration compared to WT littermates, which we interpret as an effect of increased α6* nAChR expression. Additionally, cholinergic signaling has been implicated in depressive behaviors, with smokers having higher rates of clinical depression compared to non-smokers.^31^ Drugs targeting nAChRs show antidepressant-like effects in rodent models^32^ and varenicline, a partial agonist at α4β2 and α6-containing nAChRs, can have antidepressant effects in human smokers.^33^ Our results show that varenicline reduces immobility time in the tail suspension test only in female PKCε^−/−^ mice, further demonstrating a difference in nAChR-mediated behaviors in male versus female PKCε^−/−^ mice. The mechanism by which varenicline produces antidepressant-like responses is unclear, but we speculate that the effect we observe in female PKCε^−/−^ mice may be attributed to increased α6* expression. Another potential interpretation of this result is that reduced immobility time could be attributed to enhanced locomotor activation in female knockouts, as low doses of varenicline (0.3 mg/kg) can cause increases in locomotor activity in male Wistar rats.^34^ Interestingly, a study from Mineur et al found that 1.5 mg/kg varenicline can reduce immobility time in the TST in male C57B6/J mice,^35^ while 1.0 mg/kg varenicline had no impact in the tail suspension test in male mice in our study. These different results may be attributable to differences between background strains and dosage.

### α6 nAChRs in drug consumptions and reward

α6-containing nAChRs play a major role in mediating the physiological and behavioral responses to nicotine in the VMB and on DA neuron terminals in the striatum.^36^ Mice lacking the α6 subunit fail to self-administer nicotine,^8^ and a variety of α6-targeted antagonists reduce both nicotine-induced DA release^9,37^ and nicotine self-administration in mice.^9^ Male PKCε^−/−^ mice show decreased voluntary nicotine consumption during weeks 2-4 of a 4-week 2-bottle choice paradigm, consistent with their decreased expression of α6 nAChR subunits.^15^ In contrast, we observe that female PKCε^−/−^ mice consume more nicotine during only the first week of the same paradigm compared to WT littermates, which we attribute to their increased expression of α6* nAChRs. This result is intriguing, and to the best of our knowledge there have been no studies examining the effect of α6 nAChR upregulation or functional hypersensitivity in voluntary nicotine consumption. While α4β2* nAChRs are upregulated in response to nicotine exposure,^7^ studies show that α6* nAChRs in the striatum are downregulated following prolonged nicotine exposure.^38^ Nicotine-induced downregulation of α6* nAChRs may be one mechanism that contributes to the normalization of nicotine consumption between genotypes in female WT and PKCε^−/−^ mice in weeks 2-4. Importantly, PKCε has also been shown to directly impact the function of human α4β2-containing nAChRs via phosphorylation and assist in recovery from nicotine-induced desensitization in an *in vitro* HEKtsA201 cell model,^14^ and nicotine-induced upregulation of α4* nAChRs is dependent upon generalized PKC activity.^39^ Whether PKCε interacts directly with other nAChR subunits, including α6, is unknown.

In contrast to nicotine consumption, the role of α6* nAChRs in alcohol consumption and reward is less clear. Here, we show that female PKCε^−/−^ mice consume less alcohol overall than WT littermates in a 2-bottle choice procedure, similar to prior data showing reduced alcohol consumption in male PKCε^−/−^ mice.^16^ As female PKCε^−/−^ mice have increased expression of α6 and β3 nAChR subunits while male PKCε^−/−^ mice show a decrease in transcript for these same subunits, our findings add to the already complex contribution of α6* nAChRs to alcohol consumption. Neither male nor female α6^−/−^ mice show changes in alcohol consumption in a binge drinking-in-the-dark procedure,^40^ while the α6 antagonist *N,N*-decane-1,10-diyl-*bis*-3-picolinium diiodide (bPiDI) reduces alcohol self-administration in male alcohol-preferring rats,^11^ and mice expressing a hypersensitive α6 subunit consume more alcohol in a binge drinking assay.^26^ Work from Steffensen and colleagues suggest that α6 nAChRs can reduce alcohol-induced DA release in the nucleus accumbens,^41^ which could explain why higher concentrations of alcohol result in decreased consumption in animals with higher expression of these subunits. Future studies to measure alcohol-induced DA release in female WT and PKCε^−/−^ mice will add clarity to the role of α6 nAChRs in alcohol reward.

### Sex differences in nAChR-dependent behaviors

Studies from both humans and rodent models show that sex influences nicotinic receptors and related behaviors. Clinical data indicate that women have more difficulty achieving and maintaining nicotine abstinence, and that current pharmacotherapies for nicotine cessation are less effective in women.^42^ PET imaging studies have shown sex differences in nicotinic receptor binding^43^ as well as striatal responses during nicotine consumption.^44^ Additionally, women with alcohol use disorder experience more health complications induced by alcohol consumption, and progress more quickly to alcohol dependence than men.^4^ Similar sex differences have been observed in rodent models. Female C57BL/6 mice consume more alcohol and nicotine in a two-bottle choice procedure,^22^ are less sensitive to nicotine-induced locomotor activity, and show more anxiogenic behavior in response to nicotine compared to male animals.^45^

Sex hormones can modulate the reward system, and estradiol itself can influence gene transcription in neurons.^46^ We did not monitor estrous state in our female mice, and all data was collected in freely cycling females. We observed high variability in both transcript levels and α-CTX sensitive responses in female PKCε^−/−^ mice compared to WT animals, and it is possible that circulating estradiol impacts the transcription and subsequent upregulation of α6 and β3 nAChR subunits. One interesting target of circulating estradiol are the group I metabotropic glutamate receptors, notably mGluR5, which are expressed throughout mesolimbic reward circuitry. Circulating estradiol can interact directly with mGluR5,^47^ and PKCε activity is modulated by mGluR5 activity.^48,49^ Another possibility is that PKCε ablation results in developmental adaptations that differ between males and females, as sex hormones are important mediators of brain development and organization.^50^ As our results were obtained in freely cycling females, developmental adaptations may play a more prominent role in PKCε regulation of nAChR expression compared with circulating sex hormones. Future studies will examine the mechanism through which sex influences the activity and regulation of PKCε as well as the α6 and β3 nAChR subunits, potentially through an interaction of PKCε and sex hormones.

## Conclusion

Our study reveals a previously undescribed bidirectional effect of sex on the role of PKCε in nAChR subunit expression, resulting in distinct nAChR-dependent behaviors and drug consumption. While PKCε has received attention as a potential therapeutic target for treating alcohol and nicotine dependence, our data indicate that drugs targeting PKCε may not be effective for women, highlighting the importance of including female animals in preclinical drug abuse research.

## Acknowledgements

We would like to thank Jenna Robinson, Jamie Maertens, Fayzeh El-Banna, and Cecilia Huffman for their technical assistance in support of this project. Funding provided by F31AA026782 (JKM), 4T32 DA7234-30 (JKM), R01 AA026598 (AML), UMN Foundation (AML), T32NS105640 (MCD), R01DA034696 (KW), and R01 AA027544 (KW).

## Author Contributions

JKM and AML were responsible for study concept and design. MCD acquired and analyzed the electrophysiology data. JEM contributed to the acquisition and analysis of voluntary consumption data. AML collected and analyzed data for the tail suspension test. All other data were collected, analyzed, and interpreted by JKM. JKM drafted the manuscript and MCD, KW, and AML provided critical revisions for intellectual content. All authors reviewed content and approved final version for publication.

## References

1. Dani, J. A. & Harris, R. A. Nicotine addiction and comorbidity with alcohol abuse and mental illness. Nat. Neurosci. 8, 1465–1470 (2005).

2. McKee, S. A. et al. Smoking Status as a Clinical Indicator for Alcohol Misuse in US Adults. Arch. Intern. Med. 167, 716 (2007).

3. Van Skike, C. E. et al. Critical needs in drug discovery for cessation of alcohol and nicotine polysubstance abuse. Prog. Neuropsychopharmacol. Biol. Psychiatry 65, 269–287 (2016).

4. Agabio, R., Pisanu, C., Gessa, G. L. & Franconi, F. Sex Differences in Alcohol Use Disorder. Curr. Med. Chem. 24, 2661–2670 (2016).

5. Garey, L. et al. Sex differences in smoking constructs and abstinence: The explanatory role of smoking outcome expectancies. Psychol. Addict. Behav. 32, 660–669 (2018).

6. Nagata, K. et al. Potent modulation of neuronal nicotinic acetylcholine receptor-channel by ethanol. Neurosci. Lett. 217, 189–193 (1996).

7. Albuquerque, E. X., Pereira, E. F. R., Alkondon, M. & Rogers, S. W. Mammalian nicotinic acetylcholine receptors: from structure to function. Physiol. Rev. 89, 73–120 (2009).

8. Pons, S. et al. Crucial role of alpha4 and alpha6 nicotinic acetylcholine receptor subunits from ventral tegmental area in systemic nicotine self-administration. J. Neurosci. 28, 12318–27 (2008).

9. Madsen, H. B. et al. Role of α4- and α6-containing nicotinic receptors in the acquisition and maintenance of nicotine self-administration. Addict. Biol. 20, 500–512 (2015).

10. Kamens, H. M. et al. α6β2 nicotinic acetylcholine receptors influence locomotor activity and ethanol consumption. Alcohol (2017). doi:10.1016/j.alcohol.2017.02.178

11. Srisontiyakul, J., Kastman, H. E., Krstew, E. V., Govitrapong, P. & Lawrence, A. J. The Nicotinic α6-Subunit Selective Antagonist bPiDI Reduces Alcohol Self-Administration in Alcohol-Preferring Rats. Neurochem. Res. 1–9 (2016). doi:10.1007/s11064-016-2045-3

12. Hoft, N. R. et al. SNPs in CHRNA6 and CHRNB3 are associated with alcohol consumption in a nationally representative sample. Genes, Brain Behav. 8, 631–637 (2009).

13. Culverhouse, R. C. et al. Multiple distinct CHRNB3-CHRNA6 variants are genetic risk factors for nicotine dependence in African Americans and European Americans. Addiction 109, 814–822 (2014).

14. Lee, A. M. et al. PKCε phosphorylates α4 β2 nicotinic ACh receptors and promotes recovery from desensitization. Br. J. Pharmacol. 172, 4430–4441 (2015).

15. Lee, A. M. & Messing, R. O. Protein kinase C epsilon modulates nicotine consumption and dopamine reward signals in the nucleus accumbens. Proc. Natl. Acad. Sci. U. S. A. 108, 16080–5 (2011).

16. Hodge, C. W. et al. Supersensitivity to allosteric GABA(A) receptor modulators and alcohol in mice lacking PKCε. Nat. Neurosci. 2, 997–1002 (1999).

17. Maiya, R. et al. Selective chemical genetic inhibition of protein kinase C epsilon reduces ethanol consumption in mice. Neuropharmacology 107, 40–48 (2016).

18. Blasio, A. et al. Novel Small-Molecule Inhibitors of Protein Kinase C Epsilon Reduce Ethanol Consumption in Mice. Biol. Psychiatry 0, (2017).

19. Khasar, S. G. et al. A Novel Nociceptor Signaling Pathway Revealed in Protein Kinase C ε Mutant Mice. Neuron 24, 253–260 (1999).

20. Drenan, R. M. et al. In Vivo Activation of Midbrain Dopamine Neurons via Sensitized, High-Affinity α6∗ Nicotinic Acetylcholine Receptors. Neuron 60, 123–136 (2008).

21. Crabbe, J. C., Metten, P., Ponomarev, I., Prescott, C. A. & Wahlsten, D. Effects of genetic and procedural variation on measurement of alcohol sensitivity in mouse inbred strains. Behav. Genet. 36, 536–552 (2006).

22. O’Rourke, K. Y., Touchette, J. C., Hartell, E. C., Bade, E. J. & Lee, A. M. Voluntary co-consumption of alcohol and nicotine: Effects of abstinence, intermittency, and withdrawal in mice. Neuropharmacology 109, 236–246 (2016).

23. Touchette, J. C., Maertens, J. J., Mason, M. M., O’Rourke, K. Y. & Lee, A. M. The nicotinic receptor drug sazetidine-A reduces alcohol consumption in mice without affecting concurrent nicotine consumption. Neuropharmacology 133, 63–74 (2018).

24. Arora, D. et al. Acute Cocaine Exposure Weakens GABAB Receptor-Dependent G-Protein-Gated Inwardly Rectifying K+ Signaling in Dopamine Neurons of the Ventral Tegmental Area. J. Neurosci. 31, (2011).

25. McCall, N. M., Velasco, E. M. F. de & Wickman, K. GIRK Channel Activity in Dopamine Neurons of the Ventral Tegmental Area Bidirectionally Regulates Behavioral Sensitivity to Cocaine. J. Neurosci. 39, 3600–3610 (2019).

26. Powers, M. S., Broderick, H. J., Drenan, R. M. & Chester, J. A. Nicotinic acetylcholine receptors containing α6 subunits contribute to alcohol reward-related behaviours. Genes, Brain Behav. 12, 543–553 (2013).

27. Steffensen, S. C. et al. α6 subunit-containing nicotinic receptors mediate low-dose ethanol effects on ventral tegmental area neurons and ethanol reward. Addiction Biology (2017). doi:10.1111/adb.12559

28. McIntosh, J. M. et al. Analogs of α-Conotoxin MII Are Selective for α6-Containing Nicotinic Acetylcholine Receptors. Mol. Pharmacol. 65, 944–952 (2004).

29. Morales, M. & Margolis, E. B. Ventral tegmental area: cellular heterogeneity, connectivity and behaviour. Nat. Rev. Neurosci. 18, 73–85 (2017).

30. Kamens, H. M., Hoft, N. R., Cox, R. J., Miyamoto, J. H. & Ehringer, M. A. The alpha6 nicotinic acetylcholine receptor subunit influences ethanol-induced sedation. Alcohol 46, 463–471 (2012).

31. Fergusson, D. M., Goodwin, R. D. & Horwood, L. J. Major depression and cigarette smoking: Results of a 21-year longitudinal study. Psychol. Med. 33, 1357–1367 (2003).

32. Mineur, Y. S. & Picciotto, M. R. Nicotine receptors and depression: revisiting and revising the cholinergic hypothesis. Trends Pharmacol. Sci. 31, 580–586 (2010).

33. Philip, N. S., Carpenter, L. L., Tyrka, A. R., Whiteley, L. B. & Price, L. H. Varenicline augmentation in depressed smokers: An 8-week, open-label study. J. Clin. Psychiatry 70, 1026–1031 (2009).

34. Zaniewska, M., Mccreary, A. C., Stefański, R., Przegaliński, E. & Filip, M. Effect of varenicline on the acute and repeated locomotor responses to nicotine in rats. Synapse 62, 935–939 (2008).

35. Mineur, Y. S. et al. α4β2 nicotinic acetylcholine receptor partial agonists with low intrinsic efficacy have antidepressant-like properties. Behav. Pharmacol. 22, 291–299 (2011).

36. Exley, R., Clements, M. A., Hartung, H., McIntosh, J. M. & Cragg, S. J. α6-containing nicotinic acetylcholine receptors dominate the nicotine control of dopamine neurotransmission in nucleus accumbens. Neuropsychopharmacology 33, 2158–2166 (2008).

37. Gotti, C. et al. Nicotinic Acetylcholine Receptors in the Mesolimbic Pathway: Primary Role of Ventral Tegmental Area 6 2* Receptors in Mediating Systemic Nicotine Effects on Dopamine Release, Locomotion, and Reinforcement. J. Neurosci. 30, 5311–5325 (2010).

38. Lai, A. et al. Long-term nicotine treatment decreases striatal α6* nicotinic acetylcholine receptor sites and function in mice. Mol. Pharmacol. 67, 1639–1647 (2005).

39. Wecker, L., Pollock, V. V., Pacheco, M. A. & Pastoor, T. Nicotine-induced up regulation of α4β2 neuronal nicotinic receptors is mediated by the protein kinase c-dependent phosphorylation of α4 subunits. Neuroscience 171, 12–22 (2010).

40. Guildford, M. J., Sacino, A. V. & Tapper, A. R. Modulation of ethanol reward sensitivity by nicotinic acetylcholine receptors containing the α6 subunit. Alcohol (2016). doi:10.1016/j.alcohol.2016.08.006

41. Schilaty, N. D. et al. Acute Ethanol Inhibits Dopamine Release in the Nucleus Accumbens via 6 Nicotinic Acetylcholine Receptors. J. Pharmacol. Exp. Ther. 349, 559–567 (2014).

42. Smith, P. H. et al. Sex differences in smoking cessation pharmacotherapy comparative efficacy: A network meta-analysis. Nicotine and Tobacco Research 19, 273–281 (2017).

43. Cosgrove, K. P. et al. Sex Differences in Availability of β 2 *-Nicotinic Acetylcholine Receptors in Recently Abstinent Tobacco Smokers. Arch. Gen. Psychiatry 69, 418 (2012).

44. Cosgrove, K. P. et al. Sex differences in the brain’s dopamine signature of cigarette smoking. J. Neurosci. 34, 16851–5 (2014).

45. Caldarone, B. J., King, S. L. & Picciotto, M. R. Sex differences in anxiety-like behavior and locomotor activity following chronic nicotine exposure in mice. Neurosci. Lett. 439, 187–91 (2008).

46. Tonn Eisinger, K. R., Larson, E. B., Boulware, M. I., Thomas, M. J. & Mermelstein, P. G. Membrane estrogen receptor signaling impacts the reward circuitry of the female brain to influence motivated behaviors. Steroids 133, 53–59 (2018).

47. Grove-Strawser, D., Boulware, M. I. & Mermelstein, P. G. Membrane estrogen receptors activate the metabotropic glutamate receptors mGluR5 and mGluR3 to bidirectionally regulate CREB phosphorylation in female rat striatal neurons. Neuroscience 170, 1045–1055 (2010).

48. Hermans, E. & Challiss, R. A. J. Structural, signalling and regulatory properties of the group I metabotropic glutamate receptors : prototypic family C G-protein-coupled receptors. Biochem. J 359, (2001).

49. Olive, M. F., Mcgeehan, A. & Kinder, J. The mGluR5 Antagonist 6-Methyl-2-(phenylethynyl) pyridine Decreases Ethanol Consumption via a Protein Kinase CÏμ-Dependent Mechanism. Mol. … 67, 349–355 (2005).

50. Wright, C. L., Schwarz, J. S., Dean, S. L. & McCarthy, M. M. Cellular mechanisms of estradiol-mediated sexual differentiation of the brain. Trends Endocrinol. Metab. 21, 553–561 (2010).

